# The Impact of Formic Acid Treatment on Brain Tissues for Prion Inactivation

**DOI:** 10.1101/2025.09.16.676643

**Authors:** Dalia Shaaban, Alyssa Seerley, Lilia Crew, Clairissa Kaylor, Sayre McElroy, Emma Guter, June Pounder, Andrea Grindeland Panter

## Abstract

There are significant risks in clinical, diagnostic, and research settings to those who investigate prion diseases, due to the difficult nature of inactivating prion proteins with standard decontamination methods. Formic acid treatment has been shown to be effective for decontaminating infectious prions and commonly used in biosafety practice to prevent occupational exposure. However, the impact of formic acid protocols on the morphology of tissue samples has not been adequately documented. The goal of this study is to examine morphologic effects of formic acid treatment on central nervous system tissue, using mouse model brain hemisphere tissues that exhibit varying degrees of neurodegeneration as a model. This study included normal, non-diseased wild-type tissues and a 5xFAD model, which recapitulates aspects of Alzheimer’s Disease (AD). A model exhibiting Chronic Wasting Disease (CWD), a prion disease of deer and elk, was also used to analyze the effects of formic acid on tissues with spongiform changes. Tissues from both formic acid and untreated control treatment groups were embedded in paraffin, sectioned, stained, and imaged microscopically. Anatomical regions were analyzed and evaluated quantitatively to determine the width, area, and structural integrity of the tissue between treatment groups. Our findings demonstrated that while formic acid has been previously reported to effectively inactivate prions, it compromised the morphology of mouse brain tissues. Furthermore, the effects of formic acid were not distributed equally between regions of the brain. Age did not play a role in the morphologic changes seen in the formic acid treatment group. Interestingly, the presence of neurodegeneration in the tissues did not appear to exacerbate the effects of morphological changes post-formic acid treatment. These results emphasize the need to explore alternative prion inactivation methods that ensure the safety and reliability of handling prion-infected tissues without compromising the integrity of tissues.

## 1. Introduction

Prions are infectious, misfolded proteins that lack nucleic acid and cause progressive neurodegenerative diseases in both humans and animals. They originate from a normal cellular prion protein (PrP^c^) that undergoes a posttranslational conformational change into the disease-associated form (PrP^Sc^). This transition involves the refolding of alpha-helical regions into beta-sheet-rich structures, making PrP^Sc^ highly resistant to proteases and standard inactivation methods like autoclaving, alcohol, and radiation(Prusiner, 1994, 1996, 1998; Prusiner and Hsiao, 1994; Race and Raymond, 2004; Giles *et al*., 2008; Baune *et al*., 2023; Groveman *et al*., 2024). Current methods for prion inactivation typically involve extreme conditions such as powerful acids, very strong bases, or prolonged high-temperature autoclaving protocols, all of which are arduous and may compromise tissue specimens(Brown, Wolff and Gajdusek, 1990; Sakudo *et al*., 2011; Hughson *et al*., 2016; Williams *et al*., 2019; Sakudo, 2020; Baune *et al*., 2023; Groveman *et al*., 2024).

The accumulation of PrP^Sc^ in neural tissue leads to a cascade of neurotoxic effects, including synaptic dysfunction, neuronal loss, astrocytosis, and the development of vacuoles, ultimately resulting in spongiform changes within affected brain regions^5^. These pathogenic effects are characteristic of a group of fatal neurodegenerative disorders known as transmissible spongiform encephalopathies (TSEs). In humans, TSEs include Creutzfeldt-Jakob Disease (CJD)—with sporadic, variant, genetic, and iatrogenic forms—as well as Fatal Familial Insomnia (FFI), Kuru, and Gerstmann-Sträussler-Scheinker (GSS) syndrome(Prusiner, 1994, 1998; Prusiner and Hsiao, 1994). Examples of animal forms include Bovine Spongiform Encephalopathy (BSE), scrapie, and Chronic Wasting Disease (CWD)(Goldmann, 2008). While no cure currently exists for any of these prion diseases, emerging therapeutic strategies target prion replication and propagation(Bonda *et al*., 2016).

TSEs can arise through inherited mutations in the *PRNP* gene, spontaneous misfolding events, or transmission via contaminated tissue or instruments. Clinical presentation depends on the brain region affected, and early diagnosis is limited due to the lack of specific biomarkers and effective screening tools(Sakudo *et al*., 2011). Currently, the United States Department of Agriculture (USDA) recognizes diagnosis of TSEs through post-mortem testing, using immunohistochemistry, protein immunoblot, or Enzyme-Linked Immunosorbent Assay (ELISA) detection of PrP^Sc^. Post-mortem testing clearly is not compatible with early intervention and moreover, increases the risk of exposure in clinical, surgical, and research settings(Rutala, Weber, and Society for Healthcare Epidemiology of America, 2010; Sakudo *et al*., 2011). Exposure to materials with potential prion diseases raises significant concerns due to the long incubation period between exposure and disease and the known resistance of prions to inactivation(Rutala, Weber, and Society for Healthcare Epidemiology of America, 2010; Liu, Lü and Liu, 2024). Iatrogenic transmission of human prion diseases has occurred in many instances, notably in neurosurgical procedures where inadequately decontaminated medical instruments from infected patients were reused(Blättler, 2002; Sakudo *et al*., 2011; Bonda *et al*., 2016). This has been reported in cases involving corneal grafts, dura mater transplants, and contaminated cadaver-derived pituitary hormones, previously used for growth hormone therapy, resulting in cases of prion disease(Blättler, 2002; Kobayashi, Kitamoto and Mizusawa, 2018). Additionally, transmission has occurred with stereotactic electroencephalogram (EEG) electrodes and neurosurgical equipment that was insufficiently decontaminated between procedures(Blättler, 2002). Considering these iatrogenic transmission events, the difficulty and unconventional decontamination protocols needed to inactive PrP^Sc^, and the fatal nature of TSEs, healthcare systems have adopted discard policies or enhanced containment protocols for surgical instruments used on suspected prion cases(Rutala, Weber, and Society for Healthcare Epidemiology of America, 2010; Bonda *et al*., 2016). However, with these policies come financial burdens and rigorous waste protocols. Beyond iatrogenic exposures, several documented cases involved laboratory research personnel who performed histological processing with prion-infected tissues(Brandel *et al*., 2020; Cassasus, B., 2021).

Due to the severity and highly transmissible nature of prion diseases, it is essential to avoid unintentional transmission by using appropriate inactivation methods when handling infected tissues. Previous guidelines, such as those from the World Health Organization (WHO), recommend formic acid for this purpose. While formic acid is effective in inactivating prions, there are limited publications on the effects of formic acid on tissue morphology, which raises concerns for accurate histological analysis in research experiments and clinical diagnosis. In fact, one study discusses how the use of formic acid had slightly skewed results in an investigation with sporadic CJD tissues, demonstrating the need for more investigations into the detrimental impacts of formic acid(Flønes *et al*., 2020).

Here, we report the investigation of the effects of formic acid decontamination on central nervous system (CNS) tissue morphology. Using matched brain hemispheres, one was subjected to formic acid treatment while the other served as a control. Guided by four key questions, we then explored multiple comparisons between treatments in various mouse models, including age, brain region, and disease status:

1. Does formic acid have an effect on tissue morphology? If so, is the general structure maintained, or are the tissues too damaged to do accurate assessments in histologic studies?
2. Are tissues with neurodegeneration more susceptible to damage by formic acid than normal, non-diseased tissue? If so, does specific neuropathology drive further damage by formic acid?
3. Were the morphologic changes, that were induced by formic acid treatment, equally distributed throughout the brain?
4. Does the age of the animals at harvest modulate effects of formic acid?

## 2. Materials and Methods

### 2.1 Mice

All animal procedures were conducted in compliance with the guidelines outlined in the US National Research Council’s *Guide for the Care and Use of Laboratory Animals*(*Guide for the Care and Use of Laboratory Animals: Eighth Edition*, 2011) and the US Public Health Service’s *Policy on Humane Care and Use of Laboratory Animals*. The study protocol was reviewed and approved by the Institutional Animal Care and Use Committee (IACUC) at the McLaughlin Research Institute. Mice were housed in the Animal Resource Center at the Weissman Hood Institute at Touro, McLaughlin Research Institute, a facility exclusively for mice and accredited by the American Association for Accreditation of Laboratory Animal Care.

Three different lines of mice were used for the study: C57Bl/6N, 5XFAD(Forner *et al*., 2021), and Cer-PrP-225SS (Tg1536)(Cook, Hensley-McBain and Grindeland, 2023), (Browning *et al*., 2004) (Table 1). Wild-type mice on the C57Bl/6N background strain were utilized for normal, non-diseased brain tissue comparisons between conditions, N=6 (Table 1). In addition to the wild-type mice, 5XFAD mouse model brains were included in this study to serve as a model exhibiting neurodegeneration, N=4 (Table 1). The 5xFAD (B6.Cg-Tg(APPSwFlLon,PSEN1*M146L*L286V)6799VAS/Mmjax) mouse overexpresses mutant human amyloid beta (A4) precursor protein 695 (APP) and PSEN1, a cassette containing five known genetic causes of Familial Alzheimer’s disease (FAD) in humans (APP KM670/671NL (Swedish), APP 171V (Florida), APP V7171 (London), PSEN1M146L, PSEN1 L286V); the gene cassette is controlled under a promotor which directs expression to forebrain neurons. This 5xFAD mouse line recapitulates a human neurodegenerative disease (AD), thus, displaying diseases hallmarks such as neuronal loss, amyloid plaque deposition, and gliosis(Forner *et al*., 2021). The Cer-PrP-225SS (Tg1536) expresses cervid *PRNP* at a five-fold higher level than the level of wild-type PrP expression in FVB mice. Two cohorts were included in the studies: a CWD inoculated group, and a normal brain homogenate (NBH) inoculated control group. The animals inoculated with CWD were used as a prion-positive cohort to identify the effects of formic acid in the face of spongiform degeneration (Table 1). Due to the presence of prions in these mice and biosafety concerns, no phosphate-buffered saline (PBS) controls were included in the study for this group. These mice were all at CWD endpoint ages, 9.4 months of age, N=11 (Table 1). Since the presence of neurodegeneration and spongiform change was a target of this study, the 5xFAD line and the cervidized lines did not include a young group and were included at the standard endpoints for each of their respective neurodegenerative disease manifestations.

**Table 1:** Number of brain hemisphere samples in the study shown by treatment group, genotype, and age.

| Model | Control |  | Formic Acid |  |  |
| --- | --- | --- | --- | --- | --- |
|  | Young (~4 mo) | Old (~11 mo) | Young (~4 mo) | Old (~11 mo) | Prion Endpoint (~9.4 mo) |
| WT C57B1/6N | 3 | 3 | 3 | 3 | 0 |
| 5xFAD | 0 | 4 | 0 | 4 | 0 |
| Cer-PrP-225SS (Tg1536) CWD | 0 | 0 | 0 | 0 | 5 |
| Cer-PrP-225SS (Tg1536) NBH | 0 | 0 | 0 | 0 | 6 |

### 2.2 Histology

Mouse brain hemispheres were harvested and fixed in 10% formalin (Azer Scientific, PFNBF-20) for at least 48 hours. Formic acid-treated hemi-sections were immersed in 95% formic acid (Sigma-Aldrich, 1002640100) for 1 hour with gentle agitation in a biosafety cabinet and transferred back to 10% formalin for 48 hours. The control hemi-sections were placed in fresh 10% formalin for 48 hours. Formalin was then decanted, and 1XPBS (Millipore, 6505-4L) + 0.02% Sodium Azide (Sigma-Aldrich, S8032) was added to all brain tissues. All hemispheres were prepared under standard histologic tissue dehydration and paraffin embedding conditions as referenced(Ratz-Mitchem *et al*., 2023). Standard processing includes dehydration of the tissue samples by immersing the specimens in increasing concentrations of ethanol. As paraffin is hydrophobic, the dehydration process allows for the successful embedding of the tissue. Placement in paraffin renders the samples firm enough to obtain thin sections. Sagittal sections were taken at 5 μm, mounted on positively charged slides (Avantor, 48311-703), and stained with hematoxylin (VWR, 95057-844) and eosin (VWR, 95057-848) (H&E) or a similar rapid differential stain kit (MWI Vet One, 254335699) to assess the morphology of the tissues.

### 2.3 Microscopy and Data Analysis

Images were taken on a Zeiss AxioImagerM1 microscope with either a Pixielink A623C or an Axiocam 208 (Cer-PrP-225SS brains only) color camera, morphometry analyzed using Fiji ImageJ© image-processing package and refined for publication using Photoshop 2025 (Adobe v.26.8.1). Brain structures were measured in millimeters after the settings were established in Fiji based on micrometer images from the corresponding camera. All comparisons between genotypes were made on sections representing similar medial-lateral regions of the brain to ensure uniformity. The images were taken on sections at approximately position 138 in the Allen Brain Atlas, which is 1800 μm from the medial side of the hemisphere. Cortex layer 1-6 thickness was measured in two separate measurements at a standard location immediately dorsal to the hippocampus CA1 and averaged for use in statistical analyses. The hippocampi were measured in Fiji ImageJ© using the freehand command to encircle the entire hippocampus. Measurements for each individual animal and brain region were performed twice and averaged for use in the statistical analysis.

### 2.4 Statistical Analysis

Statistical significance was determined using Student’s t-test (two-tailed) for pairwise comparisons in genotypes, unless otherwise indicated, performed in Microsoft Excel (v16.96.1) or GraphPad Prism (v10.5). Data are presented in graphs as median ± SD with statistical significance indicated as * p ≤ 0.05, ** p ≤ 0.01, ***P < 0.001, ****P < 0.0001.

## 3. Results

### 3.1 Question 1: Does formic acid have an effect on tissue morphology? Is general structure maintained?

On gross examination, all genotypes of the brain hemispheres appeared smaller after formic acid treatment, especially in length (Figure 1A,B). The image depicts a left untreated control hemisphere (Figure 1A) and the corresponding right formic acid-treated hemisphere (Figure 1B) from a wild-type mouse. Our standard H&E staining protocol proved to be challenging as some of the formic acid treated brains appeared damaged and lifted off the slide, making it more difficult, though not impossible, to obtain comparable sections (Figure 2).

**Figure 1:**
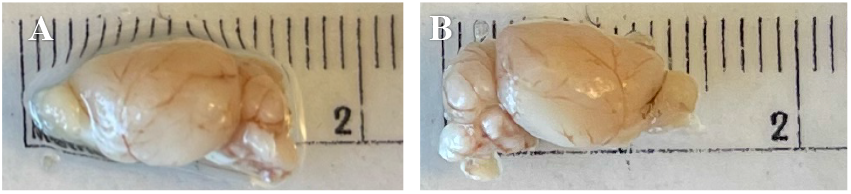
Wild-type brain hemispheres displaying smaller sizes after formic acid treatment. [A] untreated hemisphere [B] formic acid treated hemisphere.

**Figure 2:**
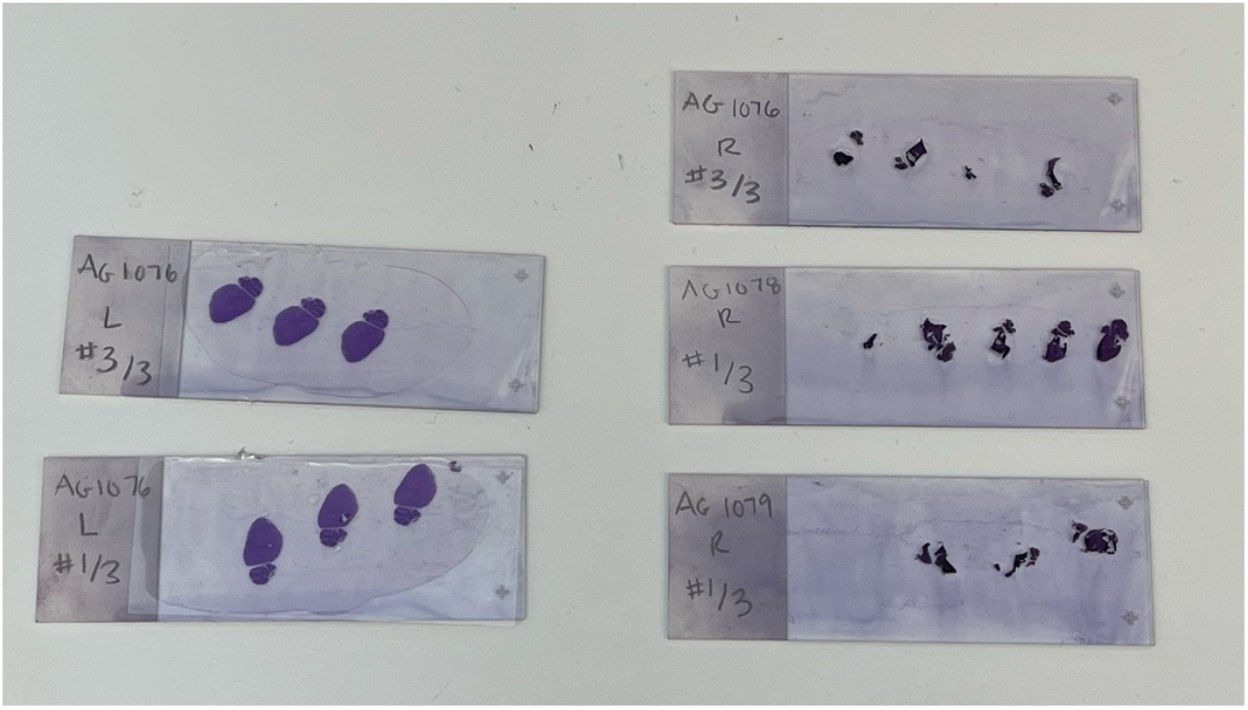
H&E-stained slides displaying detachment and damage of brain hemispheres (right) as opposed to untreated control hemispheres (left).

### 3.2 Question 2: Are tissues with neurodegeneration more susceptible to damage by formic acid than normal, non-diseased tissue?

#### 3.2.1 Formic acid treatment of wild-type and 5xFAD

The cortex width in wild-type mouse brain tissue is reduced post-formic acid treatment compared to untreated samples as shown (Figure 3). In these tissues, the average cortical width of untreated control samples measured approximately 1.0 mm, whereas formic acid-treated samples showed a reduced width of around 0.76 mm (p=0.0108) (Figure 4A). Hippocampal area in this group averaged 0.61mm^2^ in the untreated control and 0.32 mm^2^ in the formic acid treated (p=0.0013), indicating the average hippocampal area was reduced by approximately half the surface area (Figure 4A). Similar effects were observed in the 5xFAD neurodegenerative model, indicating that formic acid compromises the structural integrity of both healthy and diseased tissues (cortical width p=0.0014, hippocampal surface area p=0.0311) (Figure 4B). When comparing average loss in cortical width post-formic acid, the wild-type brains experienced 0.24 mm loss as opposed to the 5xFAD showing 0.22 mm loss, the loss of hippocampal area being 0.29 mm^2^ and 0.25 mm^2^, respectively.

**Figure 3:**
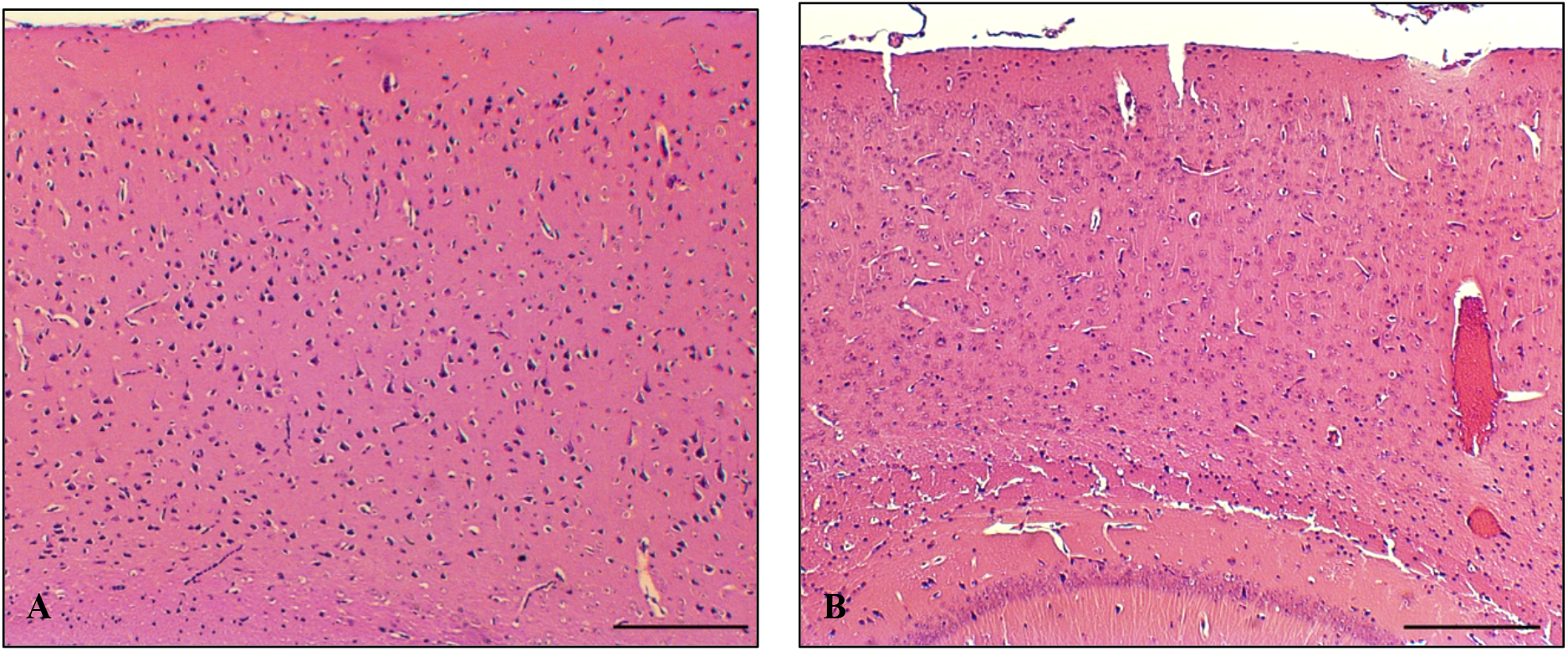
Visible cortical size differences between untreated and formic acid treated wild-type mouse brain hemispheres stained with H&E. [A] Untreated control displaying larger cortical width [B] Formic acid treated brain displaying a reduced cortex width. Scale bar: 200um.

**Figure 4:**
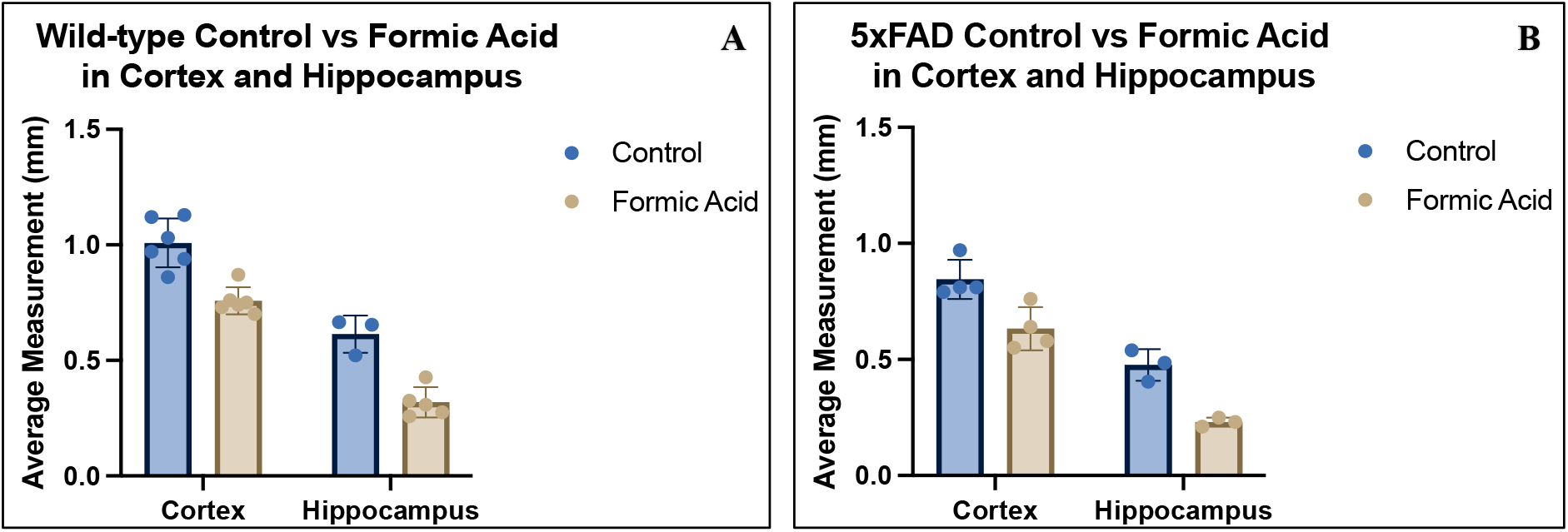
**[A]** Wild-type and **[B]** 5xFAD brain cortical width and hippocampal area measurements with control treatment shown in blue and formic acid treatment shown in beige.

#### 3.2.2 Formic acid treatment of tissues exhibiting prion disease

All Cer-PrP-225SS (Tg1536) samples were treated with formic acid due to biosafety considerations; therefore, an untreated control group was not included in this portion of the study. Samples exhibiting CWD prion disease were compared to non-diseased control tissues to determine if formic acid treatment would have a different effect on tissues exhibiting spongiform pathology. Less than a 0.01 mm change in cortical width was seen between the two groups. The average cortical CWD brain measurement was 0.38 mm wide, and NBH was 0.38 mm wide (p=0.8603), suggesting spongiform change does not influence cortical measurements post-formic acid treatment. Hippocampal area measurements had similar non-significant findings, despite the presence of major spongiform change, as shown in the representative images displaying CWD (Figure 5A) and NBH inoculated tissues (Figure 5B). Hippocampal area measurements in CWD mouse models had an average size of 0.23 mm^2^, and the NBH tissues were 0.24 mm^2^ (p=0.6983). The lack of differences between cortex width and hippocampus areas between the CWD and NBH formic acid-treated groups is interesting, as it suggests formic acid treatment does not significantly alter tissues regardless of spongiform change.

**Figure 5:**
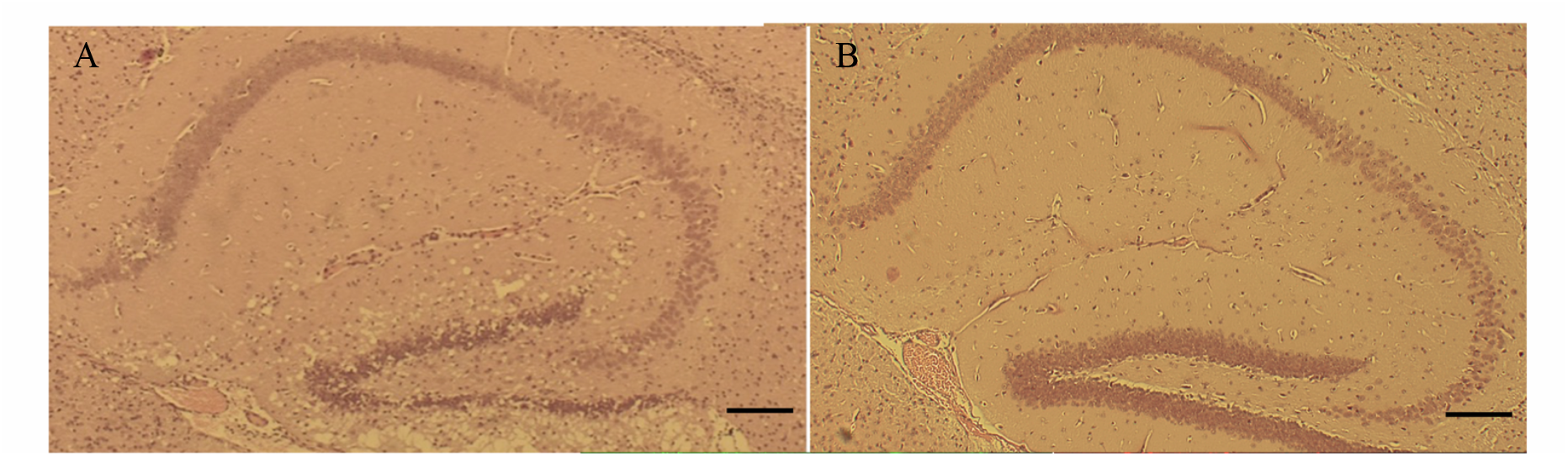
H&E-stained hippocampi in Cer-PrP-225SS mice. [A] Representative sample displaying spongiform change in endpoint CWD inoculated mice as opposed to [B] lack of spongiform change visualized in an NBH inoculated mouse. Scale bar: 200um.

### 3.3 Question 3: Were the morphological changes, that were induced by formic acid treatment, equally distributed throughout the brain?

Results indicate that the formic acid caused changes of varying degrees based on brain region. When analyzing the cortex in both the wild-type and 5xFAD mice (n=10), the average ratio of width in the formic acid treatment to control treatment was 0.82. In the hippocampus (n=6), the average ratio of surface area in the formic acid treatment to control treatment was 0.48. A two-tailed Welch’s t-test revealed a significant difference between the two groups (p= <0.0001) (Figure 6).

**Figure 6:**
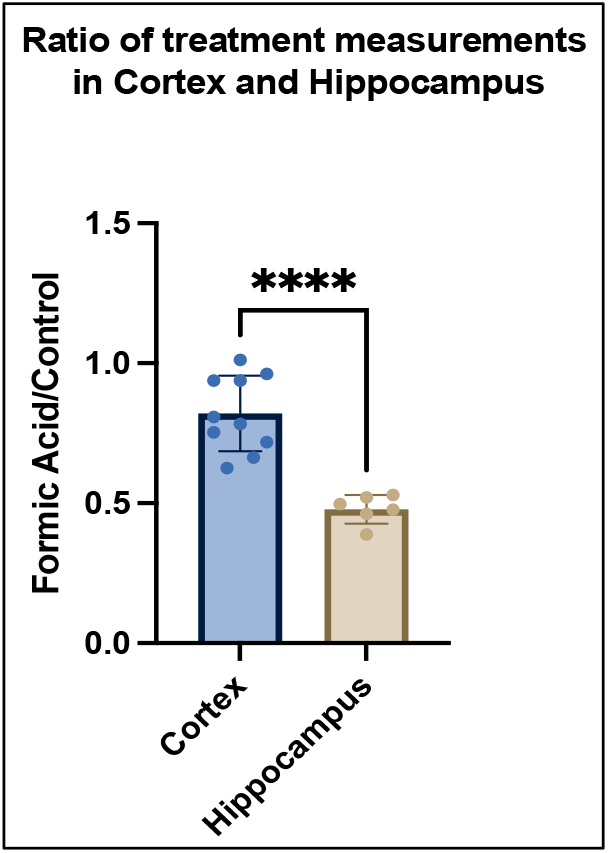
Ratios of FA treatment to control treatment measurements in both wild-type and 5xFAD mice. Cortical width measurements are indicated in blue; the hippocampal area is indicated in beige.

### 3.4 Question 4: Does the age of the animals at tissue harvest modulate effects of formic acid?

The age of mice did not appear to exacerbate the effects of formic acid treatment on the samples, as there were no significant differences when comparing the cortex measurement differences between the treatment groups in young and old mice (p=0.8388). When comparing the 3.8-month-old young group, the difference in average cortical width was 0.23 mm between the formic acid-treated samples and the untreated controls, and the 11.8-month-old group had a difference of 0.27 mm between the two treatment groups.

## 4. Discussion

Histologic assessments of CNS tissue, requiring sharp instrumentation, are often performed in both diagnostic and research settings. To ensure the safety of the personnel performing the histologic assessments, this tissue must be decontaminated appropriately with chemicals such as formic acid, which may damage the tissues or produce inconsistent results. The results of this study demonstrate a significant impact of formic acid treatment on the morphology and size of mouse brain tissue, as seen in our cortical and hippocampal measurements. These findings align with our hypothesis that formic acid, while effective in prion inactivation(Taylor *et al*., 1997), may induce morphological changes that could impact the utility of tissue samples in diagnostic and research contexts. Until alternative decontamination methods are discovered, histology professionals may be tempted to handle dangerous samples without using prion-inactivating methods due to the damaging effects of formic acid. The insurmountable cost of potential disease transmission or of lives lost highlights the danger of this sort of practice and furthers the importance of finding improved methods for prion decontamination of tissue samples(Bonda *et al*., 2016; Brandel *et al*., 2020; Cassasus, B., 2021). Our studies addressed four questions regarding the effects of formic acid treatment on brain tissue samples.

First, the results of this study demonstrate a significant impact of formic acid treatment on the morphology of mouse brain tissue in the cortical and hippocampal regions in all groups analyzed. This reveals the potential compromise of tissue integrity associated with this prion inactivation method. Most of the tissues were useful for histologic studies; however, the variability in quality and damage after long staining protocols and washing steps, will impact comparative studies and drive the use of less traditional staining protocols.

Upon staining, many of the treated brains showed partial detachment from slides, fissures, and overall structural degradation that were not observed in their sister control hemispheres. Due to the observed changes, new techniques had to be employed in some of the samples to preserve tissue integrity for analysis. A rapid differential Romanowsky stain, commonly used in pathological settings, allowed for less abrasive and efficient staining on some samples that were too fragile to undergo the normal processing steps for H&E. These two staining techniques have been shown in previous studies to be comparable when visualizing specimens in diagnostic settings(Jörundsson, Lumsden and Jacobs, 1999).

Second, we found that preexisting neurodegeneration did not appear to exacerbate the damaging effects of formic acid. The results show that formic acid compromised the morphology of both normal brain tissue and tissue with pre-existing neuropathology similarly. To determine if specific neuropathology drives further damage by formic acid, we used two different models of neurodegeneration with differing neuropathological profiles. The first model, the 5xFAD line, exhibits neuron loss, synaptic dysfunction, amyloid-beta plaque deposition, and neuroinflammation. The presence of neurodegeneration in the 5xFAD model did not appear to influence formic acid morphologic changes, as there was not a significant difference between wild-type and 5xFAD in average cortical width or hippocampal area post-formic acid.

The second neurodegenerative disease mouse model, the CWD inoculated Cer-PrP-225SS, exhibits a TSE and overt spongiform degeneration, representing a typical phenotype in which formic acid decontamination would be necessary. In this Cer-PrP-225SS (Tg1536) model, both CWD-inoculated and NBH-inoculated brains were included as a comparison between a non-diseased brain and a diseased brain treated with formic acid. We predicted that the spongiform change may render these brains more prone to shrinkage and damage in comparison to the non-diseased NBH group; however, the measurements proved to be very similar between the two groups in the face of formic acid treatment.

Third and interestingly, we observed that the hippocampus surface area after formic acid treatment was approximately half the size of the surface area measured in control samples. In the cortex, the width in samples treated with formic acid was reduced by a third when compared to the controls. These results came as a surprise due to the position of these regions in the brain. With the hippocampus being medial to the cortex, we expected that the cortex might have a greater proportion of shrinkage due to the fact that it is a larger structure and would have a higher exposure allowing it to uptake more formic acid. This implies that there may be a need for further studies on the impact of formic acid in specific brain regions, and histological assessments should consider the unequal reduction of regions in formic acid-treated samples.

Fourth and not surprisingly, our results indicate that age did not have a significant impact on cortical measurements after formic acid treatment between approximately 4- and 12-month-old wild-type mice. As the mouse brain is considered to be structurally and functionally similar to an adult by 3 months of age(Hammelrath *et al*., 2016), we were comparing two different ages of mature brains, however, the treatment of neonatal tissues may be considered in future studies.

## Conclusion

Research with prion-infected tissues presents major challenges regarding both biosafety and accurate comparable results in research studies. To avoid and prevent future exposures and incidents like the ones stated earlier, it is critical that acceptable and reliable methods of prion inactivation are established. Many compounds have been tested for prion decontamination and demonstrated varying degrees of efficacy(Race and Raymond, 2004; Giles *et al*., 2008; Rutala, Weber, and Society for Healthcare Epidemiology of America, 2010; Baune *et al*., 2023; Groveman *et al*., 2024); however, few of these have been used for specimens destined for histology(Taylor *et al*., 1997). While formic acid proves to be an effective method for prion inactivation(Taylor *et al*., 1997), its impact on brain tissue morphology, as demonstrated in this study, emphasizes the need for further investigation into alternative strategies that provide the same effectiveness as formic acid but can preserve tissue integrity. Alternative strategies would, preferable increase the reliability, reproducibility of data and the safety of handling prion-infected tissues in clinical, diagnostic, and research settings.

## Ethical Statement

The research presented here was approved by the Weissman Hood Institute at Touro University; McLaughlin Research Institutional Animal Care and Use Committee under protocol number 2023-AG-100 and biosafety protocol number 2022-AG-IBC5.

## Acknowledgements

We would like to express our gratitude to Bridget Gray for her assistance with fixation and embedding procedures. Her contributions were valuable in supporting the successful completion of this study.

